# The GABA_A_ receptor RDL modulates the auditory sensitivity of malaria mosquitoes

**DOI:** 10.1101/2024.06.05.597506

**Authors:** DA Ellis, CA Alampounti, A Suppermpool, M Georgiades, E Freeman, J Bagi, S Tytheridge, D Terrazas-Duque, JT Albert, M Andrés

## Abstract

Malaria mosquitoes mate in crowded, and noisy, swarms. A vital stage of their precopulatory behaviour involves detecting the faint flight tone of a mating partner amidst the noise from hundreds of other mosquitoes. This exquisite sensory performance is enabled by a complex auditory system with remarkable features. One such feature is their vast efferent control system, which provides the mosquito ear with the required plasticity to adapt to various external and internal changes. In this paper, we study the auditory role of GABA, one of the main efferent signalling molecules in the mosquito ear, and its interactions with octopamine, another neurotransmitter of the efferent network. We show that GABA is released in the malaria mosquito auditory nerve and that GABA receptors are expressed in the ear, including the GABA_A_ receptor Resistance to Dieldrin (RDL), a target for the evolution of insecticide resistance. Using picrotoxin to antagonize RDL receptors, we discovered multiple auditory effects of GABA at the mechanical and electrical level. At the mechanical level, blocking RDL promotes the erection of antennal fibrillae and the cessation of flagellar self-sustained oscillations. These effects are not observed in knockouts of the octopamine receptor *AgOctβ2*, suggesting that RDL auditory mechanical effects are mediated through octopamine release in the mosquito ear. Electrically, picrotoxin injection increases the spontaneous firing of auditory neurons and direct current (DC) responses of the nerve to mechanical stimulation both in wildtype and *AgOctβ2* mutants, indicating that RDL may also modulate auditory sensitivity via octopamine-independent pathways. In summary, our experiments uncover distinct auditory roles of GABA, as well as synergistic roles with octopamine. The data show a fundamental role of RDL in controlling the auditory function of malaria mosquitoes and implicate RDL signalling in mating of natural malaria mosquito populations.

## Introduction

Malaria mosquitoes have complex ears that allow them to detect the faint flight tones of mating partners in noisy swarms. The complexity of the mosquito auditory system manifests itself at different levels, including its exquisite sensitivity (1), elaborate neuroanatomy (2,3) and vast efferent control (4,5). The efferent system, which involves the release of neurotransmitters by neurons descending from the brain, has been shown to modulate mosquito auditory function (4,6). Because mosquito audition is a prerequisite for mating partner detection, the mosquito auditory efferent network also offers novel targets to disrupt mosquito reproduction (7).

The mosquito auditory efferent system releases octopamine, serotonin and gamma-aminobutyric acid (GABA) in different regions of the mosquito inner ear, or Johnston’s organ (JO) (4,5), although recent research suggest that its pharmacology could be far more complex (6). Efferent input modulates mosquito hearing at multiple levels. The complete ablation of efferent signalling triggers the onset of self-sustained oscillations (SSOs) in the male flagellum or sound receiver (8). SSOs are large spontaneous oscillations of around 350 Hz that have been suggested to act as an amplifier of female flight tones in the swarm. Moreover, release of octopamine in the ear controls the daily pattern of antennal fibrillae erection in malaria mosquitoes (6,9). Octopamine also modulates male auditory tuning in a circadian-time dependent manner which likely boosts the audibility of females in the swarm (6,10). In the dengue mosquito *Aedes aegypti*, serotonin modulates the male mosquitoes auditory tuning and affects phonotactic responses (11). The transient nature of swarming behaviour suggests that efferent input might confer auditory plasticity to mosquitoes, allowing them to adapt their auditory function to the unique sensory requirements of the swarm. Accordingly, the efferent innervation pattern is species-specific and highly sexually dimorphic, reflecting differences in the swarm auditory ecology between species and sexes (8).

Despite the extensive GABAergic innervation of the mosquito auditory nerve (4,5), GABA’s role in audition has not been studied in detail. It has been shown that GABA is released from type I efferent terminals in the auditory nerve of *Culex quinquefasciatus* mosquitoes (4,5), where it modulates the mechanical frequency tuning and nerve responses. We recently reported the transcriptome of the *Anopheles gambiae* ear (12), and found a number of GABA receptors expressed, including ionotropic (GABA_A_) and metabotropic (GABA_B_) receptors. GABA_A_ receptors form GABA-gated chloride channels, and are targeted by different insecticides (13,14). Of particular interest is the GABA_A_ receptor subunit RDL (Resistance to Dieldrin), which was first identified as the target of the insecticide dieldrin (15).

Mutations at this locus conferring resistance to dieldrin spread rapidly through wild populations of *Anopheles* as a result of widespread insecticide use in the 1950s and 1960s. Intriguingly, it has been reported that dieldrin-resistant males, specifically those with homozygous resistant mutations, have reduced mating success (16,17). However, the cause of this remains unclear. Work in the 1970s in *Anopheles stephensi* mosquitoes found that blocking RDL signalling through picrotoxin induces the erection of the antennal fibrillae in males (9). By applying different sections of the mosquito body to baths of picrotoxin, it was found that erection is dependent on RDL receptors that are located outside of the JO, most likely in the thoracic ganglion. Because the antennal fibrillae only become erect at swarm time, this effect of picrotoxin, which resembles that of octopamine (9,12), suggests a link between RDL auditory modulation and swarm time. Such a role for GABA is further supported by its connection to the circadian clock in different biological mechanisms across insects (18–21).

In this paper, we first show that GABA innervates the auditory nerve of malaria mosquitoes outside the JO and that *Rdl* is broadly expressed across JO neurons, but more intensely in auditory neurons located in distal JO regions. We use a pharmacological approach to study the auditory role of RDL by injecting picrotoxin, a well characterised antagonist of RDL receptors. We examine mechanical and electrical effects and report that RDL-mediated GABA signalling modulates the mosquito auditory physiology at multiple levels. By injecting picrotoxin, we confirm previous observations that blocking RDL induces fibrillae erection. We then use Laser-Doppler Vibrometry (LDV) to study the auditory mechanics, both in unstimulated and stimulated conditions. We find that picrotoxin causes a shift in the flagellar mechanical state towards the cessation of SSOs, and upon stimulation, a reduction in tuning sharpness. By recording from the auditory nerve, we observe that picrotoxin leads to spontaneous electrical activity and enhanced DC component responses upon mechanical stimulation, suggesting that blocking RDL reduces auditory thresholds. Moreover, picrotoxin fails to elicit mechanical changes, such as fibrillae erection, in mosquitoes that carry a knock-out mutation at the β-adrenergic-like octopamine receptor, *AgOctβ2*, indicating a functional link between GABA and octopamine to modulate the mechanics of the mosquito ear. Some of picrotoxin’s electrical effects are still present in *AgOctβ2* knockouts, suggesting that they are independent of octopamine and possibly directly mediated by the GABAergic innervation of the auditory nerve. This hints at two separate pathways - one mechanical, one electrical- of GABAergic control on the malaria mosquito auditory function.

## Materials and methods

### Mosquito rearing and entrainment

*An. gambiae* G3 mosquitoes were reared in our insectary at t he UCL Ear Institute. *AgOctβ2^-^* transgenic mosquitoes were created in a G3 genetic background (6) and maintained by intercrossing. When they were required, a 3xP3-GFP fluorescent marker was used to screen homozygotes by brightness. Every fifth generation, homozygote males would be screened and backcrossed to G3.

The colony, along with experimental individuals, were regularly genotyped by PCR. Experimental pupae were sex-separated, and males and females were hatched in different cages. Mosquitoes were then transferred to environmental incubators (Percival I-30 VL Multipurpose Plant Breeding Chamber, CLF PlantClimatics GmbH, Germany) for circadian entrainment. Entrainment conditions consisted of 12 h: 12 h light/dark cycle with 1-hour ramped light transition to simulate dawn (ZT0-ZT1, ZTX is the formalized notation of an entrained circadian cycle’s phase) and dusk (ZT12-ZT13). Light intensity during the light phase was 80 μmolm-2 s-1 from 4 fluorescent lamps. Temperature and relative humidity were kept constant at 28 °C and 80%, respectively. Mosquitoes were exposed to the circadian entrainment in incubators for three days before performing any experiment. All experiments were conducted with 3-to-7-day old mosquitoes.

### Immunohistochemistry

Following removal of the proboscis, mosquito heads were fixed in 4% paraformaldehyde for 1 h at room temperature (4,22). After fixation, heads were embedded in albumin/gelatin, post-fixed in 6% formaldehyde overnight at 4 °C and sectioned (40 μm). Sections were washed in phosphate buffered saline (PBS) 0.3% Triton X-100 and blocked in 5% normal goat serum and 2% bovine serum albumin. Primary antibodies used were monoclonal rabbit anti-glutamic acid decarboxylase (anti-GAD, 1:1000; Sigma Aldrich) and the conjugated primary antibody anti-HRP-Cy3 (1:500, Jackson ImmunoResearch, Code: 123-165-021). Secondary antibodies used correspond to Alexa Fluor Dyes (1:500; Thermo Fisher). Samples were mounted in DABCO and visualised using a Zeiss 880 confocal microscope.

### Gene expression profiles

Gene expression data for male and female JO were extracted from datasets available in (6). GABA receptor orthologs were manually extracted and normalized counts for each transcript were plotted.

### Hybridization Chain Reaction (HCR) fluorescence *in situ* hybridization

Male mosquito heads were fixed, embedded in formalin and sectioned at 40 µm slices (see Immunohistochemistry Section). HCR was performed on slices using an adapted protocol from Choi et al (2018) (23). Briefly, sections were incubated with PBS with 0.5% Triton X-100 for 1 hour at room temperature (RT). Sections were then prehybridized with prewarmed HCR hybridization buffer (Molecular Instruments, USA) for 40 minutes at 37°C. HCR probes were designed to target all splice isoforms of RDL (AGAP006028) and contain initiators that correspond with hairpins (B5) labelled with Alexa 647 fluorophore (see Supplementary Table 1). Probe solutions (Thermo Fisher Scientific, UK) were prepared with a final concentration of 24nM per HCR probe in HCR hybridization buffer.

Sections were then incubated in probe solutions overnight at 37°C. Excess probes were removed by washing 4 x 15 minutes with prewarmed probe wash buffer (Molecular Instruments, USA) at 37°C followed by 2 x 5 minutes of 5x SSCT buffer (5x sodium chloride sodium citrate and 0.1% Tween) at RT. Preamplification was performed by incubating samples with amplification buffer (Molecular Instruments, USA) for 30 minutes at RT. Hairpin h1 and hairpin h2 were prepared separately by snap cooling 4µl of 3µM stock at 95°C for 20 minutes and 20°C for 20 minutes. Sections were then incubated with h1 and h2 hairpins and anti-HRP conjugated with Alexa 488 (AB2338965, Jackson Immuno Research, UK, 1:200) in 200µL amplification buffer overnight in the dark at RT. Excess hairpins were removed by washing thoroughly the next day with 2 x 5 minutes and 3x 30 mins of SSCT at RT. Specimens were then imaged using a 20x water-dipping objective and an LSM 980 confocal microscope with Airyscan 2 (Zeiss).

### Fibrillae erection assessment

Experiments were performed at ZT4 to avoid any endogenous induction of antennal fibrillae erection that occurs at swarm time. Mosquitoes were anesthetized on ice and their thorax glued to a Teflon rod using blue-light-cured dental glue. Borosilicate microcapillaries (Drummond) were pulled with a Sutter micropipette puller and sharpened as described below. A Nanoject III was used to inject 0.2 μl of different picrotoxin concentrations (dissolved in 10% DMSO Ringer (24)) in the thorax side. Five mosquitoes were injected for each concentration and the proportion that erected their fibrillae was recorded over a 30-minute window. This experiment was repeated three times.

### Tests of auditory function

#### Laser-Doppler vibrometry (LDV)

Glass vials containing five mosquitoes were removed from incubators at ZT12 and anaesthetised on ice. They were then mounted on blue tac at the end of plastic rods using curable dental glue as previously described (6,8). The mosquito body was immobilised via glue application to minimise disturbances caused by mosquito movements but leaving thoracic spiracles free for the mosquito to breathe. The left flagellum was glued to the head and further glue was applied between the pedicels, with only the right flagellum remaining free to move.

Following this gluing procedure ^8^, the rod holding the mosquito was placed in a micromanipulator on a vibration isolation table, with the mosquito facing the Laser-Doppler vibrometer at a 90° angle. To minimise mechanical disturbances, the laser was focused at the second flagellomere from the flagellum tip in males. For electrophysiological recordings, a reference electrode was inserted in the mosquito thorax and a recording electrode was inserted in the auditory nerve. All recordings were made using a PSV-400 Laser-Doppler Vibrometer (LDV, Polytec) with an OFV-70 close up unit and a DD-500 displacement decoder. For the unstimulated free fluctuation recordings of the flagella, PSV Software was used to acquire 1-minute recordings. Spike 2 version 10 was used for data collection. All measurements were taken in a temperature-controlled room (22°C) at the different circadian times specified in each experiment.

#### Electrophysiology and electrostatic stimulation

The same mosquitoes mounted above were used for nerve recordings during electrostatic stimulation of the flagellum. Tungsten electrodes were sharpened by dipping in 10M KOH and applying a current. A reference electrode was inserted into the thorax of the mosquito and a recording electrode near the base of the right pedicel (but not into the pedicel). The reference electrode was then used to raise the electrostatic potential of the mosquito to −20 V relative to ground, enabling push/pull stimulation of the flagellum using two-electrostatic actuators placed either side of the flagellum, and fed with inverse copies of the respective command voltages. To calibrate the range of force used, the control voltage was adjusted for each mosquito such that the maximal stimulation output from our control interface resulted in 8000 nm displacement of the flagellum. During stimulation, compound action potentials were then measured using the recording electrode, and in parallel, precise measurement of flagellar displacement was measured using the LDV. All input/output was controlled and recorded using Spike 2 interfacing with a CED 1401.

#### Compound injection procedure

Picrotoxin (10 µM, Cayman Chemical) was dissolved in Ringer (24) 0.1 % DMSO (Dimethyl sulfoxide, Merck). Injections were prepared fresh on the day of the injection experiment. Sharpened micro-capillaries were filled with either picrotoxin or control Ringer solution. Borosilicate microcapillaries (World Precision Instruments) were pulled using a Sutter micropipette puller. These were sharpened with forceps by breaking the tip, then back-loaded with solution. To inject the compounds, the tip of the micro-capillary was inserted into the thorax of a mounted mosquito and the solution was injected to flood the insect with solution so as to perfuse the JO. In all injection experiments, recordings were made at two distinct stages: 1) baseline prior to any injection; 2) following injection of either 10µM picrotoxin or control solution (0.1% DMSO Ringer). This protocol allowed us to collect data before and after compound injection in the same experiment for comparative purposes, as well as ensuring that the mosquito was healthy prior to any injections.

### LDV recording and data analysis

Many of the protocols for data analysis have been published (6). For those analyses, we include a summarised version here.

#### Frequency-modulated sweep stimulation and analysis

Mosquitoes were presented with 1-second-long periods of electrostatic stimulation, interleaved by 1 second-long, free fluctuating periods, during which the flagellum does not receive any stimulus and is allowed to oscillate freely. Stimulation consisted of an electrostatic waveform as driving force with constant amplitude that swept linearly from 0 Hz to 1000 Hz (forward sweep), or 1000 Hz to 0 Hz (backward sweep). Forward sweeps were presented followed by no stimulus, then backward sweeps followed by no stimulus and so on, repeating 25 times. Each of these units of stimulation and rest is referred to as a run. All data have undergone a curation process to ensure that the laser was appropriately positioned on the mosquito flagellum during an experimental run. This was done through a diagnostic channel which measures and records the laser backscatter of the LDV. The use of both forward and backward sweeps allows us to account for any hysteresis-driven latency by averaging.

##### Unstimulated free-fluctuation analysis

We used the unstimulated data prior to the stimulated counterpart to determine the flagellar mechanical state, which is determined by looking at the amplitude distribution of the flagellar motion. Details of the analytical pipeline have been published (6). Briefly, the amplitude distribution of a fully quiescent flagellum follows a standard normal distribution, whereas an SSOing animal would instead follow an arcsine distribution. An added complication comes from the appearance of transient mechanical states which tend to lie somewhere in between a QUIES and SSO state. To account for those states, we generate a mixed distribution comprised of an arcsine and normal distribution, controlled by a weight parameter α which ranges between 0 (denoting purely QUIES) and 1 (denoting purely SSO). To permit some level of flexibility on the mechanical state, the cut-off condition for QUIES and SSO animals is set between [0 – 0.1] and [0.9 – 1.0] respectively and require a goodness of fit to account for 99.7% of the data. We further formalise the SSO definition by adding another criterion where an animal is only classified as SSO if it sustained its oscillatory behaviour for three consecutive experimental runs, corresponding to a total span of six seconds. This condition was set to avoid attributing the SSO label to a briefly oscillating animal. Analysis was performed using Mathematica v.13.1.

Fitting procedures complicate the calculation of velocity amplitudes of free-fluctuations. We used fast Fourier transform analysis of free-fluctuation data to extract the frequency and velocity of peak responses.

##### Analysis of unstimulated nerve

These analyses used only the unstimulated portions of each run and all analyses were performed in Python (https://github.com/mosquitome/sweep-n-sleep). For each individual, data was analysed in discontinuous 1-second windows. Data was median subtracted to centre it around zero, and a peak-calling algorithm (find_peaks, SciPy) was used to find all peaks in the negative sign. These were stringently filtered to only include peaks more than 5 standard deviations from zero. Inter-peak intervals were calculated within these 1 second windows. For the generation of random peak trains, data was simulated for an equal number of individuals as real data, each with the same number of 1 second windows, with each window containing the same number of peaks as real data. The time assigned to each peak, however, was randomly chosen from a uniform distribution (random_choice, NumPy) between 0 and 1 seconds at a sampling rate of 20k.

##### Frequency-modulated sweep stimulation analysis

These analyses used only the stimulated portion of each run. For spline-fitting analyses, stimulus, nerve and laser data were averaged over the 25 repeats for each individual. Nerve data was then split into low and high frequency components in Spike 2 by applying a DC remove (Time constant =0.01 s) and smooth (Time constant =0.01 s) filter, respectively. The rest of the analyses were performed using Python. The frequency of the stimulus at any given time was interpolated (poly1d, NumPy) from a linear fit (polyfit, NumPy) through a spectrogram of the stimulus (spectrogram, SciPy). The envelope of the laser data was determined by downsampling by a factor of 5, then finding minima and maxima throughout the signal. Quadratic splines were then fit to the lower and upper envelopes (UnivariateSpline, SciPy), incrementally increasing the smoothing factor until only one peak was present above the 99^th^ percentile (find_peaks, SciPy). The average of lower and upper peaks was used for analyses. The envelope of the AC component of the nerve was determined using rolling minimum and maximum windows of 100 data points (rolling, Pandas). Splines were fit as for laser data using a threshold of the 90^th^ percentile. The position of the lower peak was used for analyses (with its corresponding value in the upper spline used to determine the peak-to-peak amplitude). A single spline was fit directly to the DC component of the nerve using the same approach as for the AC component.

For damped harmonic oscillator fitting, we used the same approach as in (6). The profile of the stimulated data follows a well-known line shape that is defined by the driven damped harmonic oscillator. The so-called ‘envelope’ of the waveform is extracted using a statistic-sensitive nonlinear iterative peak clipping (SNIP) method of background estimation. The envelope is then fitted through the theoretical expectation from the driven harmonic oscillator and different fitted parameters are extracted to describe the biophysics of the system including: *F*_0_the driving force, m the effective mass of the flagellum, *ω*_0_the natural frequency of the oscillator, the peak frequency or frequency at which the oscillatory system achieves its maximum amplitude as a function of the driving frequency, *ζ* the damping ratio and *ω* the driving frequency. A commonly used derived parameter that describes a damped oscillator is the Q-factor, defined as *Q* = ½*ζ*, which can be used to qualitatively interpret the level of dampening experienced by the flagellum. Analysis was performed using Mathematica v.13.1.

### Statistical analysis

Samples sizes for LDV experiments were determined based on published data on Dipteran antennal LDV measurements ^8,87,88^. Within-group variation estimates were calculated as part of standard statistical tests and were reasonable for the type of recordings. Statistical tests for normality (Shapiro-Wilk test with a significance level *of p < 0.05*) were used for each LDV dataset. These were generally found to be non-normally distributed; thus, median and median absolute deviation values are reported throughout. For the fibrillae erection experiments and amitraz exposure experiments, Fisher’s exact test (FET) was used to compare the frequencies of fibrillae erection and mechanical state across the different compound injections.

For the free fluctuation and unstimulated nerve measurements, paired sample Wilcoxon tests or paired t-test were used to perform pairwise comparisons between before and after compound injections. For frequency-modulated sweep stimulation, some samples had to be omitted, particularly after injection, as we were unable to perform robust spline-fitting on averaged data. This led to uneven sample sizes in a number of groups when comparing before with after compound injection, reducing the power of paired tests. Welch’s independent t-tests or Mann-Whitney tests were therefore used for these comparisons, depending on the distribution of the compared data.

Statistical tests were performed in R 4.2.2 or Python 3.9.

## Results

### GABA innervates the auditory nerve of male malaria mosquitoes

The mosquito ear is an antennal hearing organ composed of the sound receiver, or flagellum, and the mosquito’s “inner ear”, the Johnston’s organ (JO, Fig. 1A). To analyse the pattern of GABAergic innervation of the auditory system on *An. gambiae*, we immunostained antennal sections with an anti-GAD antibody (glutamic acid decarboxylase, which converts glutamate into GABA and is commonly used as marker of GABAergic neurons (25)). This antibody extensively labelled the auditory nerve outside the JO (Fig. 1B), corresponding to the type I terminals of the auditory efferent system (5) as previously described in *Cx. quinquefasciatus* mosquitoes (4).

**Fig 1:**
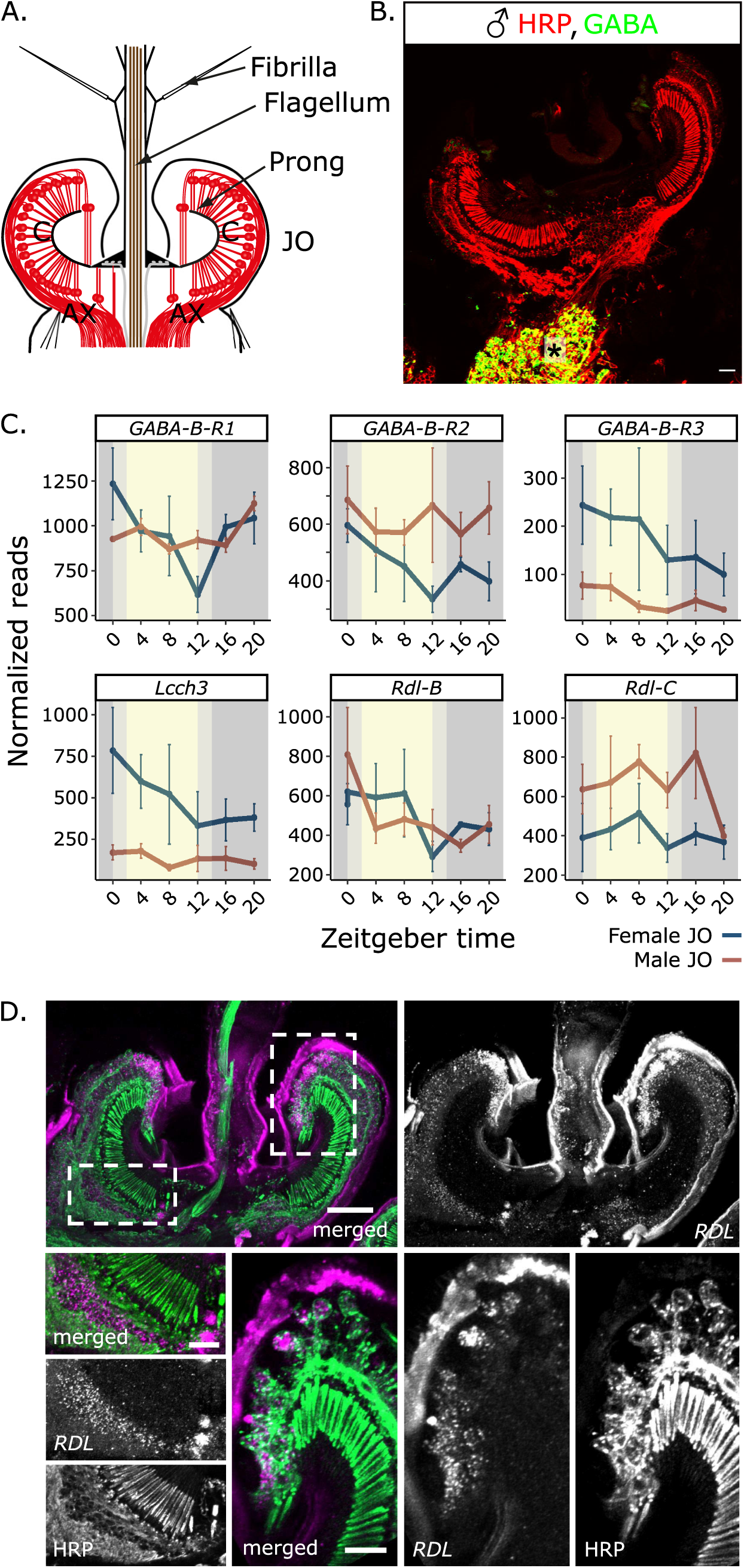
GABA in the auditory system of malaria mosquitoes. **A**) Schematic of a male mosquito JO depicting the sound receiver (flagellum) and the auditory organ, the Johnston’s organ (JO), located at its base in the second antennal segment. **B**) Male JO section labelled with an anti-GAD antibody in green (glutamic acid decarboxylase, an enzyme implicated in GABA synthesis) showing GABAergic innervation of the auditory nerve (asterisk), and co-stained with the neuronal marker anti-HRP (red). **C)** Expression of different GABA receptors in male and female JO throughout the day (ZT12 represents dusk or laboratory swarm time). Normalized read numbers (mean ± se), data have been extracted from (6). **D**) Hybridization chain reaction-fluorescence *in situ* hybridization (HCR-FISH) of JO section to detect *Rdl* transcripts (magenta) and counterstained with anti-HRP (green). *Rdl* transcripts are found broadly distributed across auditory neuron cell bodies (bottom left panels) but accumulate in the most distal neuronal cell bodies (bottom right panel), probably including neurons belonging to type B and distal type A scolopidia. Scale bars: 25 µm, zoom-ins: 10 µm. Sample size = 4 male mosquitoes.

Next, we investigated GABA receptor expression in transcriptomic data of the malaria mosquito JO (6). GABA receptors can be subdivided into two types: GABA_A_, which are ionotropic GABA-gated ion channels, and GABA_B_, which are metabotropic G-protein coupled receptors (26,27). Two GABA_A_ receptor genes (*Rdl* and *AGAP000038*, potential orthologue of ligand-gated chloride channel homolog 3, Lcch3) and three GABA_B_ receptor genes (GABAB-R1, GABAB-R2 and GABAB-R3) were expressed in the malaria mosquito JO (Fig. 1C), with some showing clear levels of sexual dimorphism. This suggests complex GABAergic modulation of the mosquito auditory pathways. Amongst the most highly expressed receptors were two distinct splice isoforms of the insecticide target, RDL.

To analyse the distribution of *Rdl* expressing neurons, we performed *in situ* hybridization chain reaction (FISH-HCR) against both *Rdl-B* and *Rdl-C* in sections of male JOs. Positive labelling was broadly observed in neuronal cell bodies (Fig. 1D), suggesting that *Rdl* is widely expressed across auditory neurons. Some labelling could be observed in clusters outside of cell bodies. Although this probably corresponds to neuronal regions outside the cell body, we cannot rule out that *Rdl* is also expressed by support cells. Labelling was more intense in neuronal cell bodies located in the more distal region of the JO (Fig. 1D), in the same area as type B scolopidia (2). Moreover, cell bodies belonging to the most distal type A scolopidia, which bind to the tip of prongs, were also intensely labelled. These results suggest differential *Rdl* expression across JO neurons.

### RDL signalling modulates different mechanical properties of unstimulated malaria mosquito ears

In malaria mosquito males, the sound receiver, or antennal flagellum, is covered by small fibrillae that follow a rhythmic pattern of erection under circadian clock control and become extended at swarm time (9). Picrotoxin, a noncompetitive antagonist of GABA receptors formed by RDL homomers (28), has been shown to induce fibrillae erection in *An. stephensi* mosquitoes (9). We tested the effects of injecting picrotoxin on the fibrillae erection in *An. gambiae* mosquitoes. We performed the experiments at ZT4, during the laboratory day, when the fibrillae are normally collapsed. We found that injecting increasing levels of picrotoxin led to an increasing proportion of individuals with erected fibrillae. Upon 100 µM picrotoxin injection, 87% ± 19 (mean ± standard deviation) of mosquitoes had fully erect fibrillae, with the remaining 13% ± 19 showing partial erection (Fig. 2A). Whilst injecting ≥ 100 µM picrotoxin led to a maximal response in this assay, these high doses had off-target effects, even causing visible twitching in some mosquitoes. We therefore injected 10 µM – approximately the EC_50_ dose – for all experiments herein.

**Figure 2:**
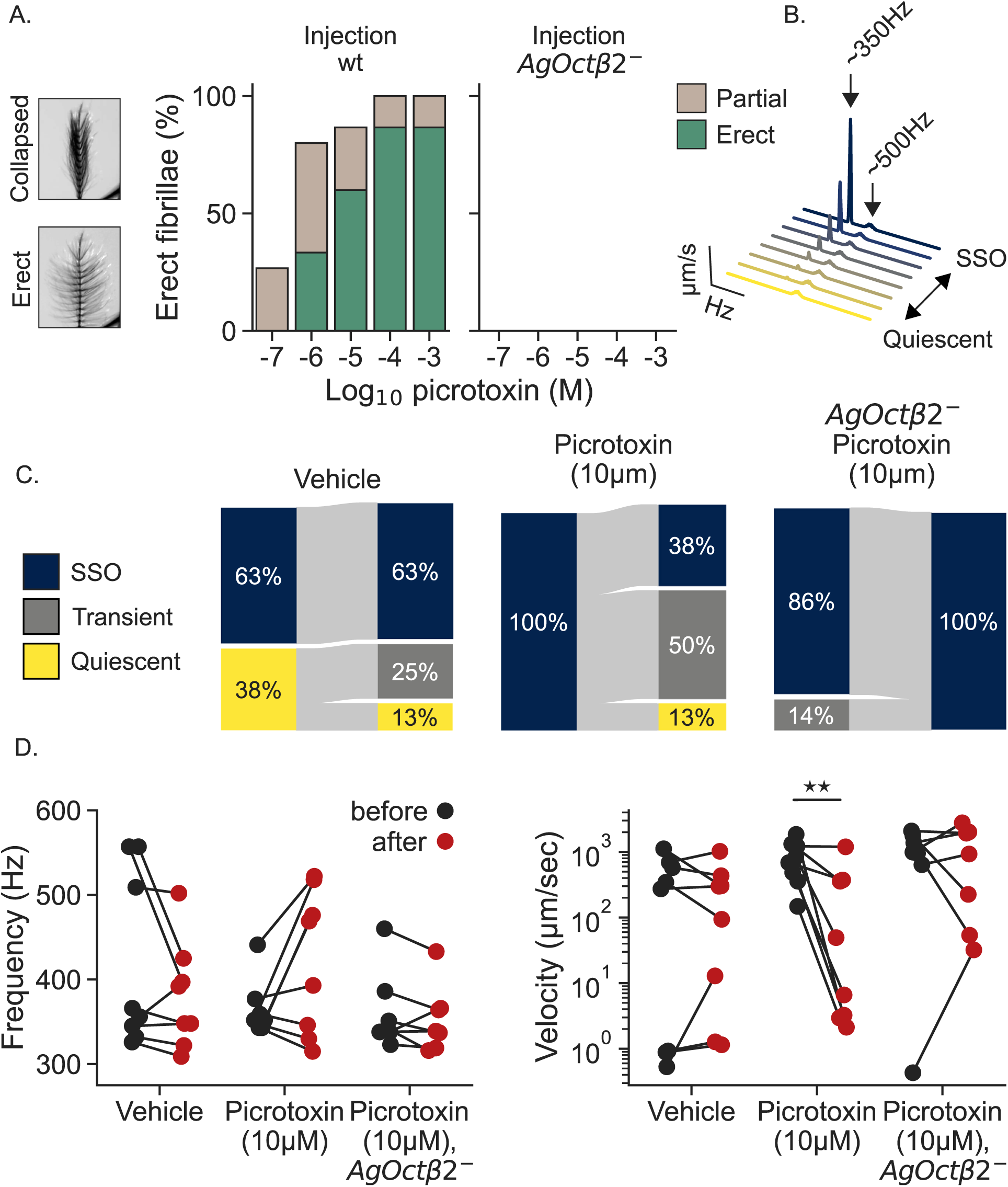
Picrotoxin induces fibrillae erection and SSO cessation in unstimulated malaria mosquito ears. **A)** Effects of picrotoxin injection on fibrillae erection were observed at ZT4, when fibrillae are naturally collapsed. Picrotoxin induces fibrillae erection in wt, but not *AgOctβ2* mutants. Left: example image of an antenna in a collapsed and erect state. Centre: Percentage of wt mosquitoes displaying partially (brown) or completely (green) erect fibrillae in a 30-minute window after picrotoxin injection. Bars show the mean of 3 replicates, with 5 mosquitoes injected for each concentration, per replicate. Right: Percentage of *AgOctβ2* mutants displaying partially or completely erect fibrillae in a 30-minute window after picrotoxin injection (there were none). **B)** Illustration of how mosquito flagella transition between two distinct mechanical states in unstimulated conditions. These states are characterized by their frequency and amplitude. SSO flagella show large oscillations of ∼ 350 Hz, quiescent flagella oscillate at ∼500 Hz with much smaller amplitudes and broader frequency tuning. **C)** Mechanical states of the mosquito flagellum were analysed at ZT12 after control and 10 µM picrotoxin injection. No changes were observed in the distribution of mechanical states upon control injection, and the proportion of SSOing mosquitoes did not change. However, 10 µM picrotoxin injection caused a shift from SSO to transient or quiescent states in 63% of mosquitoes (Fisher’s exact test, p-value < 0.05). In *AgOctβ2*^-^ mosquitoes, there were no changes in this direction. **D)** Effects of picrotoxin on the frequency and velocity magnitude of free fluctuation recordings. The frequency did not significantly change after picrotoxin injection in wildtype animals (although some mosquitoes showed a clear increase). However, the velocity magnitudes of the oscillations decreased upon picrotoxin injection (Paired t-test, p < 0.01). Control injection, or picrotoxin injection in *AgOctβ2^-^,* did not cause any changes in these auditory parameters. Significant differences between injection effects starred **p <0.01. Sample sizes: control = 8, picrotoxin = 8, *AgOctβ2^-^* picrotoxin =7.

The flagellar mechanics reflect the sum of all components mechanically coupled to the flagellum. The flagellum itself presents spontaneous activity that emerges from active energy injection by auditory neurons (8) and male mosquito flagella have been reported to present two distinct mechanical states (Fig. 2B). Some receivers show large spontaneous “self-sustained oscillations” (SSOs) of ∼ 350 Hz and velocity magnitudes of ∼1mm/s. Others, present quiescent states with best frequencies ∼ 500 Hz and much lower velocity magnitudes (∼1 µm/s). Some flagella present a transient state between both (6). Although the role of the different mechanical states in nature is not fully understood, SSOs seem to provide a mechanism for males to amplify female flight tones (8,29,30). A laser Doppler Vibrometer (LDV) directed to the tip of the flagellum was used to measure the frequency and amplitude of its motions and thereby infer the flagellum’s mechanical state (see Methods (6)).

We compared the flagellar mechanical states before and after injecting 10 µM picrotoxin or control (0.1% DMSO) solutions. It is worth noting that the factors influencing the mechanical state of the mosquito flagellum are still unknown and variation in mechanical state exists even at baseline. However, we were interested in whether compound injection causes changes in the mechanical state, reflecting changes in mechanical components of the auditory system. Control injections did not affect the mechanical state of the mosquito flagella; there was only a minor shift from quiescent to transient states, and the proportion of mosquitoes showing SSOs did not change (Fig.2C, control_before_: 63% SSO, 38% quiescent; control_after_: 63% SSO, 13% quiescent, 25 % transient, Fisher’s Exact Test p > 0.05). By contrast, we observed that 10 µM picrotoxin injection caused a cessation of SSOs in 50% of mosquitoes and a shift towards transient or quiescent states (picrotoxin_before_: 100% SSO; picrotoxin_after_: 38% SSO, 13% quiescent, 50% transient, Fisher’s Exact Test p < 0.05).

We also extracted the oscillator frequency and velocity from free fluctuation recordings (Fig. 2D). Overall, the oscillator frequency did not change after picrotoxin injection, although some of the picrotoxin-injected animals showed a clear increase in contrast to control-injected (Fig. 2D). However, the velocity of the oscillations robustly decreased after picrotoxin injection, changing by an order of magnitude (Fig. 2D, picrotoxin_before_: 760 ± 431 µm/s showing median ± median absolute deviation (mad); picrotoxin_after_: 28 ± 26 µm/s, paired Wilcoxon signed-rank test p < 0.01). In summary, parameters describing the amplificatory state of the ear shift towards quiescence after picrotoxin injection.

### RDL-dependent mechanical control of unstimulated receivers is mediated through octopamine

The mechanical effects of picrotoxin are similar to those of octopamine, which we recently found to modulate different auditory parameters via the *AgOctβ2* receptor (6). Among these, *AgOctβ2* mutants fail to erect their fibrillae either at swarm time, or upon octopamine exposure, which causes fibrillae erection in wildtype mosquitoes. To analyse potential functional links between RDL and octopamine, we performed picrotoxin injections in knockout mosquitoes of *AgOctβ2*. Our results showed that none of the mechanical effects that we observed after picrotoxin injection in wildtype mosquitoes were observed in *AgOctβ2^-^*knockouts. Mutant animals did not exhibit fibrillae erection with picrotoxin, even at the highest concentrations (Fig. 2A). Moreover, *AgOctβ2-* mosquitoes did not change their mechanical state upon picrotoxin injection, remaining in, or shifting to a SSO state upon picrotoxin injection (Fig. 2C). Furthermore, the velocity of spontaneous vibrations, which decreased in wildtype mosquitoes upon injection, did not change in mutant animals (Fig. 2D). Together, these results suggest a connection between AgOctβ2 and RDL to modulate the mechanics of the malaria mosquito ear and fit a model where RDL inhibits downstream octopamine release, which in-turn, binds AgOctβ2 and induces mechanical changes in the ear.

### Blocking RDL leads to an increase in the spontaneous activity of the auditory nerve

By analysing the spontaneous activity of the malaria mosquito ear, we have shown that RDL modulates the ear’s mechanical properties via AgOctβ2. Because we observe extensive GABAergic innervation of the auditory nerve (Fig. 1B), we next investigated potential effects of picrotoxin in the electrical activity of the JO using electrophysiology. Empirically, one of the most striking results of picrotoxin injection was a visible increase in spontaneous nerve activity (Fig. 3A). Our stimulation protocol alternates between 1 s of sweep stimulation and 1 s without stimulation. To quantify the spontaneous activity, we extracted compound nerve activity recordings during the periods without stimulation and called spiking events in this data (Fig. 3B; see methods). Note that these “peaks” do not necessarily correspond to single-fiber activity as our nerve recordings are extracellular and record activity from numerous axons. We use the term peak to refer to depolarizing events in the auditory nerve. We calculated the number of peaks per second and observed an increase upon picrotoxin injection (Fig. 3C, peaks per s; picrotoxin_before_: 0.15 ± 0.13Hz; picrotoxin_after_: 6.80 ± 5.85 Hz, paired Wilcoxon signed-rank test p < 0.01). No change in peak number was observed upon control injection. To understand the distribution of peaks in time, and whether they showed any regularity, we calculated inter-peak intervals (within each discontinuous 1 s bin; see methods). Note that peaks quantified before picrotoxin injection are based on very few events and may represent noise, rather than true biological signal. To generate a null distribution where events were spread uniformly in time, we simulated data with the same number of individuals, runs and peaks as our real post-injection data. The inter-peak interval of real events was more heavily skewed towards zero than random events, with an exponential distribution characteristic of bursting. These bursts of neuronal firing could, for example, be caused by neuronal connectivity through e.g. electrical synapses.

**Figure 3:**
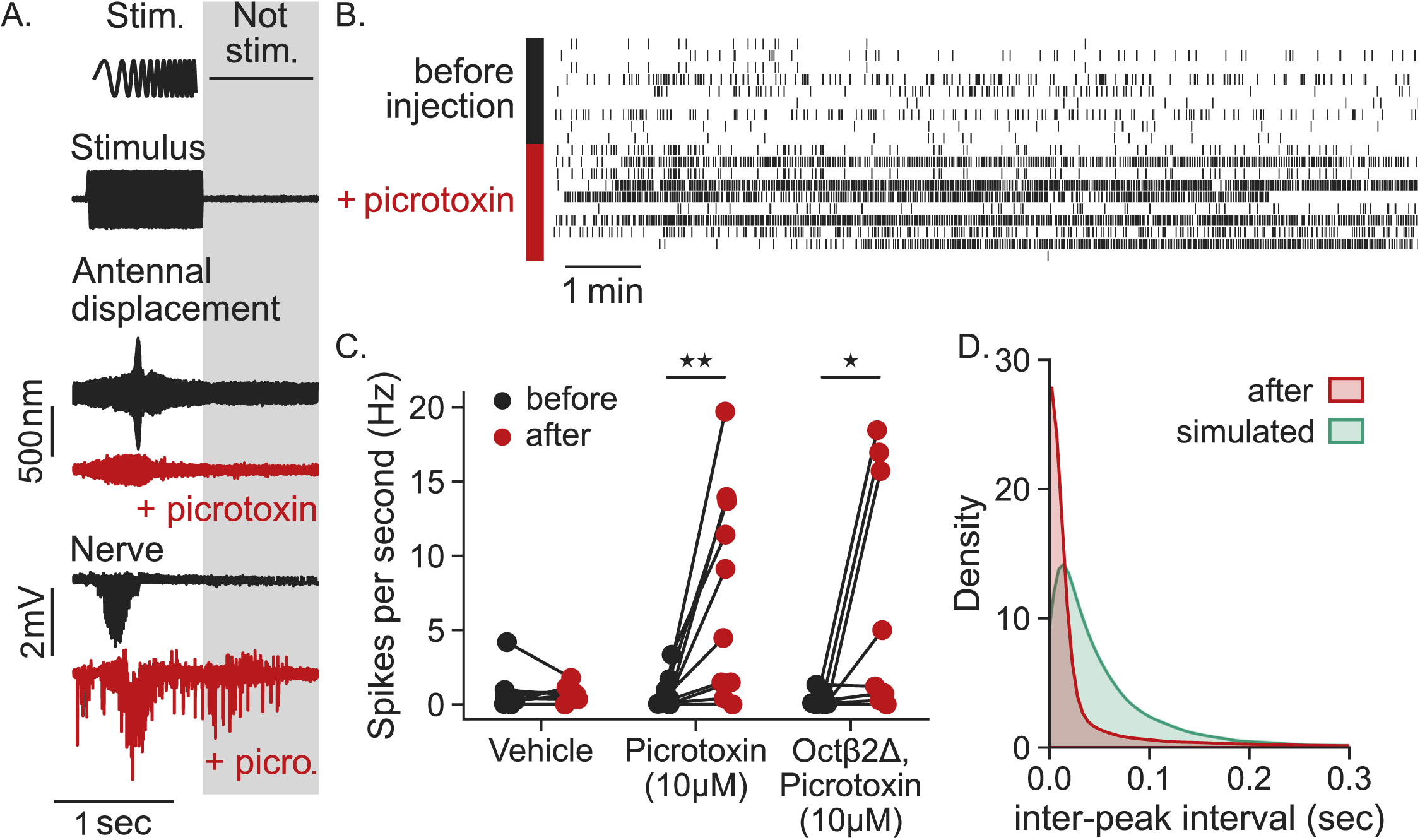
Picrotoxin increases the spontaneous activity of the auditory nerve. **A)** Example traces showing experimental paradigm, including stimulus (top), mechanical antennal displacement (centre) and nerve responses (bottom). Baseline response shown in black, and the response to picrotoxin shown in red. Data is from a single run of a recording from a single mosquito. Our stimulation protocol alternated between stimulation (“Stim.”) and free-fluctuating periods (“Not stim.”; grey). Data in this figure is extracted from the unstimulated portion highlighted in grey, when no stimulus was provided. **B)** Raster plot showing electrical peak incidences across the entire experiment (unstimulated sections of each run only). Each row corresponds to a single mosquito. Data shows wt mosquitoes injected with picrotoxin; other treatments are summarised in part C. **C)** Peaks per second in each condition. Each pair of points correspond to a single mosquito before (black) and after (red) injection. Injecting picrotoxin caused a large increase in the frequency of peaks (peaks/s) both in wildtype (paired Wilcoxon signed-rank test, p< 0.01) and *AgOctβ2* knockouts (paired Wilcoxon signed-rank test, p< 0.05), but not upon control injection. **D)** Distribution of the inter-peak intervals for wt mosquitoes after injection with picrotoxin (red), as well as simulations where data from after injection had peak times randomised (green). Intervals were calculated within runs. Simulated data had the same number of mosquitoes, runs and peaks within each run as after-injection mosquitoes. Only data from wt mosquitoes are shown (mutants showed a similar distribution).

### Blocking RDL increases the flagellar tuning sharpness and enhances DC component nerve responses upon mechanical stimulation

Our results show that picrotoxin modulates the spontaneous activity of the malaria mosquito ear both at the mechanical and electrical level. We further wanted to analyse whether mechanically evoked responses were also altered. To test this, we measured flagellar displacements using a laser Doppler Vibrometer (LDV) and compound action potential (CAP) responses of the nerve using electrophysiology (paradigm and example response in Fig. 3A). We used frequency-modulated sweeps (upchirps and downchirps, 0–1000 Hz) to mechanically stimulate the flagellum.

To analyse mechanical responses, we first fit splines to the envelope of averaged flagellar responses (Fig. 4A, right; see methods). By calling peaks in these splines, we could identify the amplitude of peak-to-peak flagellar responses, and the stimulus frequency that elicited them. There was atendency to an apparent decrease in peak-to-peak flagellar displacement upon picrotoxin injections that was not significant (Fig. 4A, peak-to-peak amplitude; picrotoxin_before_: 891 ± 245 nm; picrotoxin_after_: 558 ± 305 nm, Mann-Whitney p > 0.05). The stimulus frequency that caused the greatest flagellar displacement did not change after picrotoxin injection (Fig. 4A, stimulus frequency, picrotoxin_before_: 363 ± 12 Hz; picrotoxin_after_: 356 ± 37 Hz, Mann-Whitney p > 0.05).

**Figure 4:**
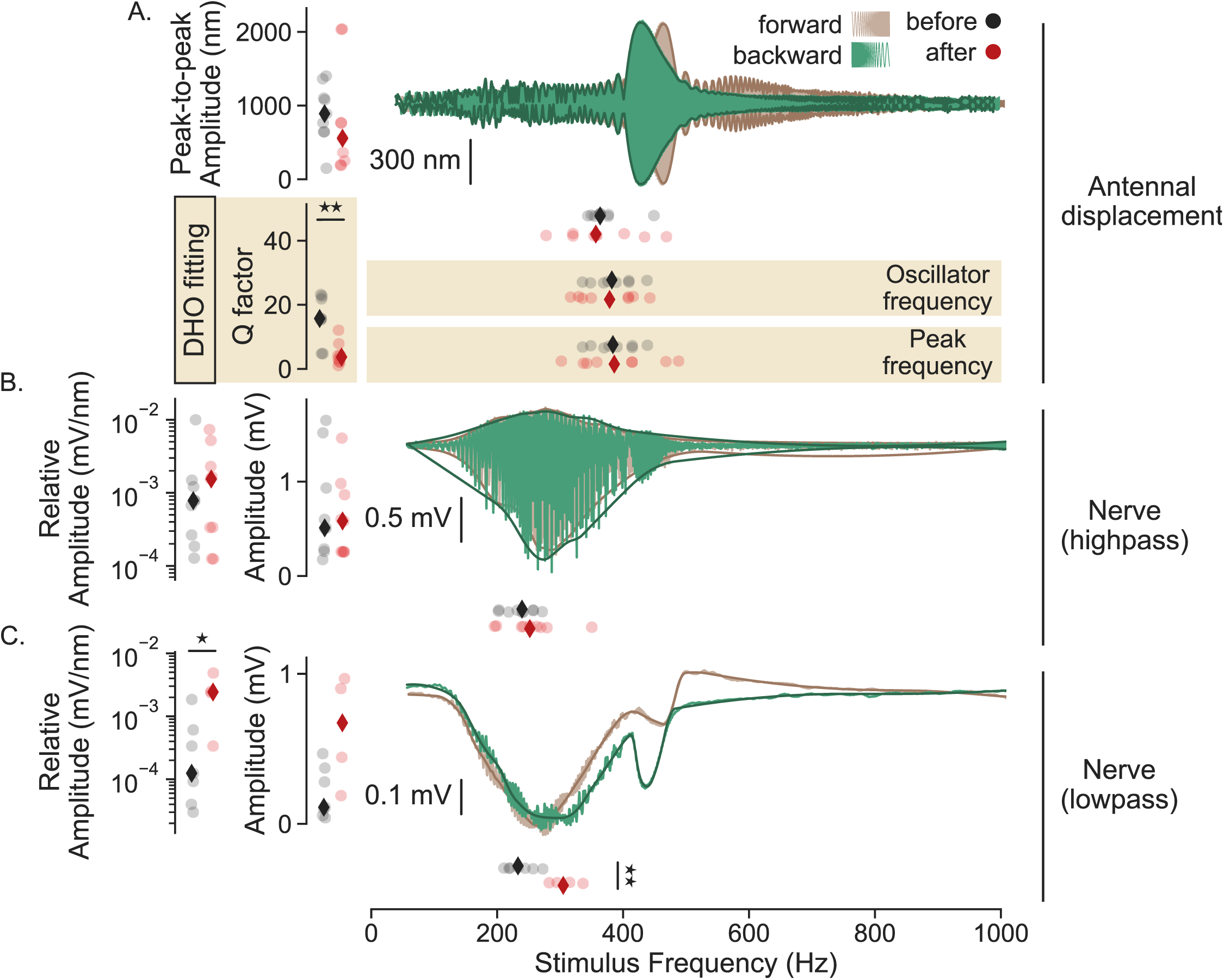
Picrotoxin increases DC component responses to mechanical stimulation. Wildtype male mosquitoes were exposed to frequency-modulated sweep stimulation. Measurements were performed at swarm time (ZT12) before and after 10 µM picrotoxin injection. Line plots (green and brown) show averaged data from one example mosquito, detailing the response to forward (brown) and backward (green) sweep stimulation at the antenna (A) and nerve (B and C). Data have been projected into the frequency domain, with the x-axis corresponding to the stimulus frequency, and the y-axis to the response amplitude. Lines with deeper hue show the splines that were fit to either the envelope (A and B) or raw data (C). Strip plots (black and red) show summary statistics extracted from these fits (average of forward and backward responses; see methods). For strip plots, each circular point represents data from a single mosquito; diamonds with deeper hue show the median. Nerve amplitude data (left panels) is shown both relative to mechanical displacement (mV/nm) and un-adjusted (mV). Relative data is shown on a log axis. **A)** Envelopes of antennal responses were either fit with splines (top, white background), or a Damped Harmonic Oscillator (DHO) function (bottom, ochre background). See methods for details of fitting procedures. For splines (top, white background): left strip plots show the peak-to-peak amplitude of flagellar displacements between upper and lower splines. Data extracted from individuals before (black) and after injection (red). Lower right strip plots show the stimulus frequency at the peak response. For DHO fits (bottom, ochre background): left strip plots show the Q factor (i.e. tuning sharpness) extracted from the fits, and right strip plots show the oscillator (top) and peak frequency (bottom; see (6)). Picrotoxin injection caused a reduction in Q factor (Welch’s independent t test, see methods; p < 0.01), but oscillator and peak frequencies did not change. **B)** AC component extracted from applying a highpass filter to compound action potential (CAP) responses. Left strip plots show spline-to-spline amplitude at the peak of the lower spline. Lower right strip plots show the stimulus frequency at the peak response. Neither the amplitude nor the frequency of the stimulus causing peak AC component responses changed after picrotoxin injection. **C)** DC component extracted by applying a lowpass filter to compound action potential (CAP) responses. All strip plots are the same as for high-pass filtered data. Picrotoxin injection causes an increase of the relative DC component amplitude (peak DC component amplitude normalized to the peak-to-peak flagellar displacement, Mann-Whitney, p < 0.05). Picrotoxin also increased the stimulus frequency at which the peak DC response occurred (Welch’s t test, p< 0.01). Significant differences between before and after injection starred *p <0.05; **p <0.01. DHO: damped harmonic oscillator. Sample size = 9, some samples had to be omitted, particularly after injection, as we were unable to perform robust spline-fitting on averaged data.

To further characterise changes in the flagellar mechanics, we fit forced damped harmonic oscillator (DHO) models to the flagellar displacements in order to extract other relevant biophysical parameters (Fig. 4A, DHO fitting). This analysis showed that injecting 10 µM picrotoxin did not affect the auditory tuning of the organ (Fig. 4A, DHO fitting) as both peak and oscillator frequencies have similar values before and after picrotoxin injection (Fig. 4A, peak frequency; picrotoxin_before_: 384 ± 30 Hz; picrotoxin_after_: 386 ± 47 Hz, Welch’s t test p > 0.05). However, injecting picrotoxin led to a significant decrease in the tuning sharpness (Q-factor) of the flagellum, indicating that the flagellum was more responsive to a broader range of frequencies (Fig. 4A, Q-factor; picrotoxin_before_: 15.7 ± 6.6; picrotoxin_after_: 3.7 ± 1.5, Welch’s t test p < 0.01). The tuning sharpness did not change upon control injections in wildtype mosquitoes (Supp. Fig. 1).

We then analysed nerve responses to sweep stimuli. We separated the alternating current (AC) and direct current (DC) components of CAP responses by applying lowpass and highpass filters (see Methods) and fit splines. This allowed us to extract the peak amplitudes, and the stimulus frequency at this peak, for both components (Fig. 4B-C). Our analysis found no significant change in the peak amplitude of the AC component after picrotoxin injection (Fig. 4B, picrotoxin_before_: 0.51 ± 0.25 mV; picrotoxin_after_: 0.58 ± 0.32 mV, Welch’s t test p > 0.05), nor the stimulus frequency causing these peaks (Fig. 4B, picrotoxin_before_: 239 ± 18 Hz; picrotoxin_after_: 252 ± 17 Hz, Welch’s t test p > 0.05).

Because picrotoxin-dependent changes in the mechanics may influence nerve responses, we also normalised nerve amplitudes to flagellar displacement amplitudes (Fig. 4B, left). However, this didn’t reveal any effect (picrotoxin_before_: 7.82e-4 ± 5.2e-4 mV/nm; picrotoxin_after_: 1.55e-3 ± 1.21e-3 mV/nm, Mann-Whitney p > 0.05).

For the DC component however, whilst the effect of picrotoxin on un-normalised amplitudes was non-significant, albeit large (Fig. 4C, picrotoxin_before_: 0.11 ± 0.07mV; picrotoxin_after_: 0.67 ± 0.26 mV, p > 0.05), the relative amplitudes (normalizing DC component amplitude to flagellar displacements) were significantly increased after injection (Fig 4C, left; picrotoxin_before_: 1.25e-4 ± 9.45e-5 mV/nm; picrotoxin_after_: 2.43e-3 ± 1.08e-3 mV/nm, Mann-Whitney p < 0.05). This suggests that blocking RDL enhances DC component responses of the nerve, independent of any effect on the auditory mechanics. The stimulus frequency causing peak responses also significantly increased after picrotoxin treatment (Fig 4C, picrotoxin_before_: 233.04 ± 14.58 Hz; picrotoxin_after_: 304.91 ± 16.15 Hz, Welch’s t test p < 0.01). Control injections did not affect DC component responses (Supp. Fig. 1).

### Picrotoxin effects on the auditory nerve are partially independent of octopamine

As the mechanical effects of picrotoxin on unstimulated ears were dependent on octopaminergic signalling, we then tested whether this was also the case for its electrical effects. We first examined how blocking RDL affected the spontaneous activity of the auditory nerve in *AgOctβ2* knockout males. In a similar manner to wildtype, injecting picrotoxin led to an increase in the number of peaks per second (Fig. 3C, peaks per s; AgOctβ2-picrotoxin_before_: 0.07 ± 0.06 Hz; AgOctβ2-picrotoxin_after_: 1.22 ± 1.22 Hz, paired Wilcoxon signed-rank test p < 0.05). Under unstimulated conditions, picrotoxin increases the spontaneous nerve activity of both wildtype and *AgOctβ2* knockout males.

We then applied the auditory stimulation protocol (Fig 3A) to *AgOctβ2^-^*mosquitoes (Fig. 5). Mechanical responses to frequency-modulated sweep stimulation showed no significant differences upon picrotoxin injection in any of the parameters tested (Fig. 5A). This supports our findings from unstimulated flagella that the mechanical effects of picrotoxin are mediated through AgOctβ2-dependent octopaminergic signalling.

**Figure 5:**
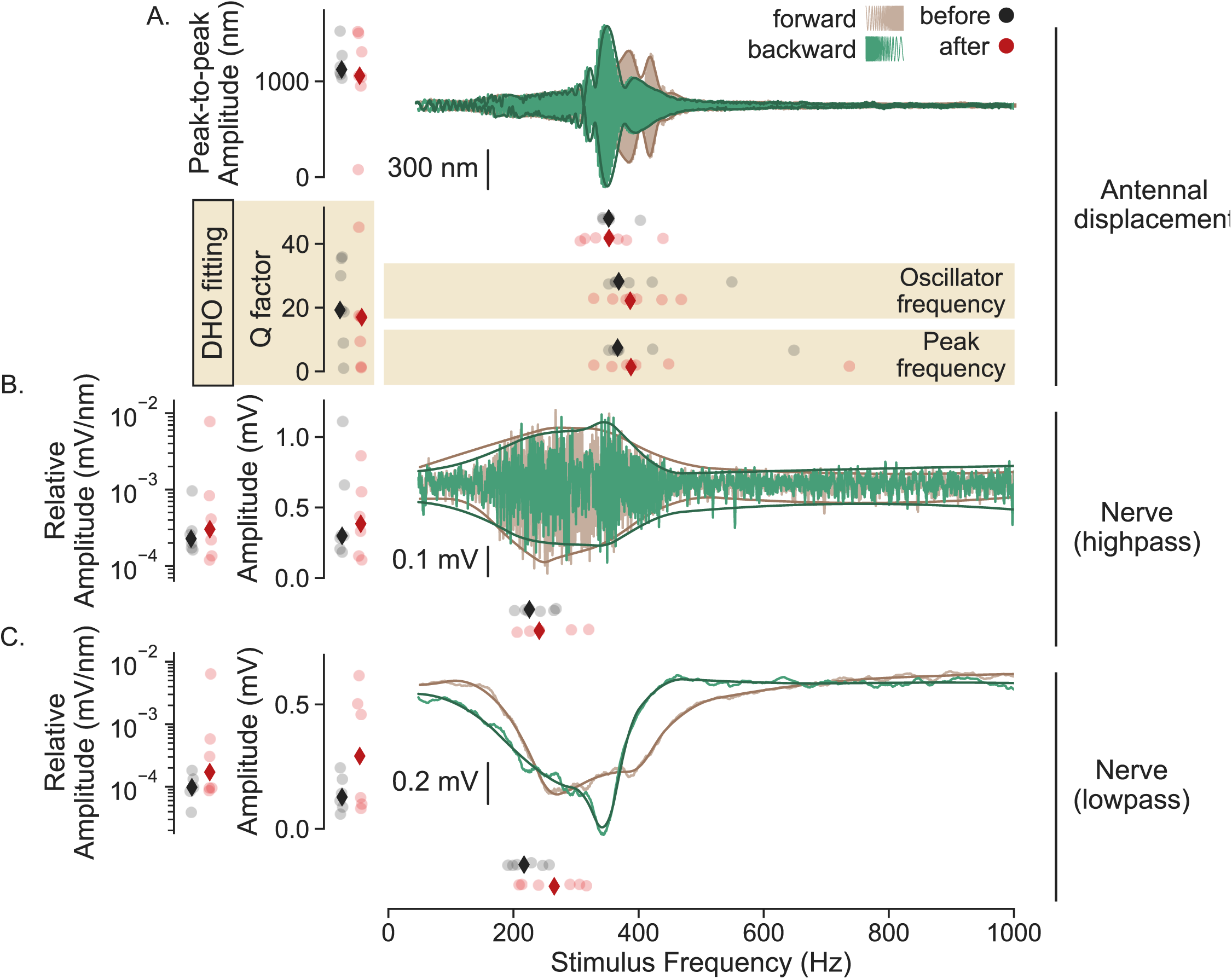
Picrotoxin effects on stimulated auditory responses in mutant *AgOctβ2^-^* mosquitoes. Male *AgOctβ2* mutant mosquitoes were exposed to frequency-modulated sweep stimulation. The data is displayed in the same way as for wildtype animals in Fig. 4 (see its legend for a detailed description of data shown). **A)** Envelopes of antennal responses were either fit with splines (top, white background), or a Damped Harmonic Oscillator (DHO) function (bottom, ochre background). See methods for details of fitting procedures. For splines (top, white background): left strip plots show the peak-to-peak amplitude of flagellar displacements between upper and lower splines. Data extracted from individuals before (black) and after injection (red). Lower right strip plots show the stimulus frequency at the peak response. For DHO fits (bottom, ochre background): left strip plots show the Q factor (i.e. tuning sharpness) extracted from the fits, and right strip plots show the oscillator (top) and peak frequency (bottom) – another set of parameters that can be extracted from DHO fits (see (6)). Picrotoxin injection did not cause any changes in the auditory mechanics. **B)** AC component extracted by applying a highpass filter to compound action potential (CAP) responses. Left strip plots show spline-to-spline amplitude at the peak of the lower spline. Lower right strip plots show the stimulus frequency at the peak response. The frequency of the stimulus causing peak AC component responses did not change after picrotoxin injection. **C)** DC component extracted by applying a lowpass filter to compound action potential (CAP) responses. All strip plots are the same as for high-pass filtered data. Picrotoxin injection causes an increase of the relative DC component amplitude, but this was not significant (Welch’s t test, p> 0.05). DHO: damped harmonic oscillator. Sample size = 9, some samples had to be omitted, particularly after injection, as we were unable to perform robust spline-fitting on averaged data.

We then assessed nerve responses in *AgOctβ2* mutants during stimulation. In contrast to the spontaneous nerve activity of unstimulated ears, we found that picrotoxin had no significant effects on nerve responses. As in wildtype animals, AC component responses did not change after picrotoxin injection, neither for absolute peak amplitudes (Fig. 5B, AgOctβ2-picrotoxin_before_: 0.30 ± 0.09 mV; AgOctβ2-picrotoxin_after_: 0.38 ± 0.23 mV, Mann-Whitney p > 0.05), nor peak amplitudes relative to flagellar displacements (Fig. 5B, AgOctβ2-picrotoxin_before_: 2.29e-4 ± 5.85e-5 mV/nm; AgOctβ2-picrotoxin_after_: 3.17e-4 ± 1.89e-4 mV/nm, Mann-Whitney p > 0.05). For DC component responses, although non-significant, we still observed an upward shift in the absolute peak amplitude in *AgOctβ2* mutants (Fig. 5C, AgOctβ2-picrotoxin_before_: 0.14 ± 0.07 mV/nm; AgOctβ2-picrotoxin_after_: 0.32 ± 0.25 mV, Welch’s t test p > 0.05). This difference was reduced when calculated relative to the flagellar displacements (Fig. 5C, AgOctβ2-picrotoxin_before_: 0.017 ± 0.008 mV/nm; AgOctβ2-picrotoxin_after_: 0.31 ± 0.24 mV/nm, p > 0.05).

These results indicate that although picrotoxin increased DC component responses in wildtype mosquitoes upon mechanical stimulation, this effect was not observed in *AgOctβ2^-^* mosquitoes. Taken together, recordings of nerve activity under stimulated and unstimulated responses suggest that picrotoxin effects on the nerve are independent of octopamine action, as shown by the changes in spontaneous activity, but can be modulated by it.

## Discussion

In this manuscript, we investigated the auditory role of the GABA_A_ receptor RDL in malaria mosquitoes.We first provided immunohistochemical evidence of extensive GABAergic innervation in the auditory nerve and identified five GABA receptors expressed in the mosquito ear, including RDL, which was amongst the most highly expressed. Because of its high level of expression and public health relevance, we then went on to subject the RDL receptor to more detailed functional analyses. *In situ* hybridization chain reaction (HCR) showed differential *Rdl* expression across JO neurons, with higher levels in neurons located in the most distal JO region, suggesting functional differences across neuronal subtypes. Using a pharmacological approach, we then blocked RDL signalling using picrotoxin, a non-competitive antagonist (or blocker) of GABA receptors formed by RDL subunits. We found that blocking RDL activity had manifold effects on mosquito auditory mechanics: it induced erection of antennal fibrillae; shifted the oscillatory state of the male flagellum from SSO to quiescent; and caused a decrease in the tuning sharpness. The fact that these effects were abolished in mosquitoes that carry knockout mutations of the β-adrenergic-like octopamine receptor, *AgOctβ2*, an octopamine receptor previously shown to modulate auditory function in malaria mosquitos (6), suggests that the mechanical effects of RDL are mediated through octopamine release in the ear. Moreover, our results indicate that GABAergic input in the auditory nerve plays a fundamental role in modulating electrical sensitivity, as picrotoxin injections affected nerve responses by increasing spontaneous activity and enhancing DC component responses to mechanical stimulation. Together, these results suggest that altering RDL signalling could severely affect the auditory performance of malaria mosquitoes.

Immunostaining with an anti-GAD antibody (glutamic acid decarboxylase, marker of GABAergic neurons) showed extensive GABAergic innervation of the auditory nerve (Fig. 1A). Because the GABAergic innervation targets the auditory nerve before the first synapsis of the auditory pathway in the antennal mechanosensory and motor center (AMMC) (11), these connections should be axo-axonic. GABAergic innervation of the auditory nerve has also been found in *Cx. quinquefasciatus* mosquitoes (4), where ultrastructural analysis showed that the connections between the auditory axons and GABAergic neurons were via axo-axonic synapsis. Although around 25% of synapses in the insect brain are estimated to be axo-axonic (31), their roles are still not well understood. Axo-axonic connections contribute to processing sensory information in different systems. In mammals, GABAergic interneurons provide modality-specific filtering of sensory information (32). In *Drosophila*, axo-axonic connections have been found in the lateral horn between projection neurons that might be involved in odour discrimination (33). They are also involved in olfactory memory formation in the *Drosophila* mushroom body (34). Moreover, axo-axonic GABAergic synapses have been shown to be highly plastic and to switch their polarity from depolarizing to inhibitory in the developing mouse brain (35).

To identify the GABA receptors mediating its auditory role, we extracted data from a recently published transcriptomic analysis of the mosquito JO (6). We identified two ionotropic (GABA_A_) and three metabotropic (GABA_B_) receptors expressed in the mosquito ear (6), suggesting both fast (GABA_A_) and slow (GABA_B_) processing of auditory information (36). From these, we selected RDL for further examination as it was the most highly expressed GABA receptor in our dataset. Moreover, RDL is the target of different insecticides that are used for mosquito control, so understanding RDL auditory function could have important implications for insecticide development and management. FISH-HCR showed positive punctae labelling *Rdl* mRNA broadly distributed across neuronal cell bodies. *Rdl* transcripts accumulated in the cell bodies of most distal auditory neurons. These results suggest RDL-mediated functional specializations of the auditory neurons within the malaria mosquito JO. In particular, cell bodies of auditory neurons forming B scolopidia, which account for only 3% of scolopidia in the JO and attach to the apical side of the prongs emerging from the flagellar base (2) (as opposed to A scolopidia, which make up approximately 97% of scolopidial units and attach to the most distal basal side of prongs) showed high *Rdl* expression. Some of the most distal neurons belonging to A scolopidia also presented high *Rdl* expression levels. These results suggest that *Rdl* is expressed by most auditory neurons, but it is most highly expressed in distal neurons. No labelling could be observed in the auditory nerve suggesting that RDL is synthesised in cell bodies to then reach its final localization in the axons.

Our pharmacological approach to block RDL signalling using picrotoxin revealed functional roles of the receptor at multiple levels of audition. Firstly, picrotoxin injection induced the erection of the antennal fibrillae. This resembled the effects that were recently described for octopamine (6). In this previous study, octopamine was found to promote fibrillae erection through the receptor *AgOctβ2* (beta 2 adrenergic-like receptor), and when this receptor was mutated, fibrillae no longer became erect. Interestingly, in this paper we have found that picrotoxin did not cause any change in the antenna of *AgOctβ2* mutant mosquitoes. This suggests that GABA acts upstream of octopamine to control its release for fibrillae erection. Fibrillar erection is strongly circadian (9); GABA has been extensively documented as acting as an output of the circadian clock (19–21,37,38). We propose that during the day, GABA inhibits octopamine release. This GABAergic inhibition would cease during swarm time to promote octopamine release in the ear and induce fibrillae erection. A study from the 1970s found that in *An. stephensi* both octopamine and picrotoxin induced antennal fibrillae erection when injected into the mosquito thorax (9). In this study, local application of octopamine in isolated antenna, but not picrotoxin, also induced fibrillae erection. Combined with our findings, these results suggest that GABAergic and octopaminergic neurons could connect at the level of the thoracic ganglion. Although the origin of the octopaminergic neurons innervating the mosquito ear is yet unknown, the anatomy of the octopaminergic system is well conserved across insects. There are a small number of octopaminergic neurons that innervate most brain regions and peripheral organs, including dorsal unpaired median (DUM) and ventral unpaired median (VUM), which originate in the subesophageal ganglion and thoracic ganglia (39–45). In cockroaches, octopaminergic DUM neurons in the last abdominal ganglion have been shown to receive GABA input and are inhibited by picrotoxin (46). Therefore, we propose that RDL expressed in octopaminergic neurons, potentially in the thoracic ganglion, constitutively inhibits octopamine release, except for the swarm period, when GABA inhibition would temporarily stop to allow octopamine release and subsequent fibrillae erection through signalling via AgOctβ2. Further anatomical and functional studies are required to test the validity of this hypothesis.

Some of the auditory mechanical effects previously described for octopamine, which included an increase in the mechanical tuning frequency (6), were not observed with picrotoxin. We interpret these differences as being a consequence of different compound concentrations in both studies.

While in the previous study, 1 mM or 10 mM octopamine injections were used to analyse their auditory effects, we used 10 µM picrotoxin injections. It is plausible that the octopamine released in the ear upon 10 µM picrotoxin injections is lower than the levels reached by 1 or 10 mM octopamine injections, so the effects we observed are milder. However, we decided to use 10 µM picrotoxin as it represented the EC_50_ dose of the fibrillae erection experiments (Fig. 2A) and higher concentrations caused visible twitching of mosquitoes, and too dramatic nerve effects that were difficult to analyse.

Apart from the mechanical role of RDL mediated through octopamine release, the extensive GABAergic innervation of the auditory nerve hinted at further functions. Indeed, picrotoxin injection had clear effects on nerve responses, including a strong enhancement of DC component responses upon mechanical stimulation. Effects of picrotoxin on the DC component of nerve responses have been previously observed in *Cx.quinquefasciatus* mosquitoes (4). Furthermore, analysing compound spontaneous nerve activity showed a large increase in spike number upon picrotoxin injection, suggesting modulatory effects on auditory sensitivity, which is one of the main roles described for auditory efferent control in vertebrates (47). As RDL is a GABA-gated chloride channel, its conductance might contribute to hyperpolarize the nerve and modulate auditory thresholds.

Injecting picrotoxin would block RDL, therefore contributing to depolarizing axons and increasing the nerve spontaneous activity. Similar effects of picrotoxin as a convulsant has been described in mice (48). The enhanced DC component of CAP responses could emerge from the summation of synaptic activity caused by the reduction in auditory thresholds induced by picrotoxin. Additionally, because we observed decreased tuning sharpness of the mechanical responses, it is plausible that the modulation of auditory thresholds contributes to frequency discrimination, as has been previously described for GABA currents in other sensory modalities (33).

It is worth noting that our study is limited by the fact that our measurements of the nerve activity are CAPs, which capture nerve activity from many neurons. The data can therefore be difficult to interpret. RDL is widely expressed in the insect central nervous system mediating GABA inhibitory signals (49). We cannot rule out that some of the increase in the nerve spontaneous activity that we observe upon picrotoxin injection originates in non-auditory pathways. However, the increased DC component responses upon frequency-modulated sweep stimulation provides evidence that RDL is directly involved in nerve responses to auditory stimulation. Moreover, our data – and others (50) – suggest the existence of auditory neuron subpopulations that might be affected differentially by picrotoxin. With the level of resolution of our electrophysiological data, these questions are difficult to solve. Single-cell electrophysiological recordings would provide the level of resolution required to interpret our results more accurately and describe RDL kinetics in single neurons.

The effects of picrotoxin on spontaneous nerve activity were also found in *AgOctβ2* mutant mosquitoes (Fig. 3C). However, effects on DC component responses to mechanical stimulation were not present (not significant), although there was a trend showing similar responses as in wildtype animals (Fig. 5C). Combining these findings with our knowledge of the anatomy of the GABAergic and octopaminergic innervation in the mosquito ear, we propose that the nerve effects of picrotoxin are independent of octopamine. This agrees with the anatomical evidence that shows that the octopaminergic fibres innervate the base of the auditory cilia, while GABA terminals locate at the auditory nerve (4). However, as picrotoxin nerve effects are milder in *AgOctβ2* mutants – no changes were observed upon mechanical stimulation – it seems that octopamine influences nerve responses to a certain degree. It is plausible that octopamine, which is released at the base of the auditory cilia, amplifies mechanical responses. Mutant animals would lack this mechanical amplification, therefore reducing nerve responses and picrotoxin effects. Further detailed investigation of different components of the efferent system on auditory thresholds are required to investigate this hypothesis.

Based on the results presented in this paper, we propose that RDL-mediated GABAergic signalling modulates malaria mosquito audition through two pathways (Fig. 6): 1) RDL expressed in octopaminergic neurons, potentially in the thoracic ganglion, inhibits octopamine release in the mosquito ear except at swarm time to influence the mechanics of the ear in a circadian-time dependent manner (e.g. fibrillae erection pattern), 2) RDL expressed in the JO auditory neurons and localized to the nerve, controls the auditory sensitivity by modulating the nerve auditory thresholds. Indeed, both pathways might be connected to modulate the auditory physiology of the malaria mosquito in the swarm. Reducing RDL-mediated GABA signalling during swarm time – likely under circadian clock control – would lead to 1) release of octopamine in the ear to induce fibrillae erection and cause other mechanical effects, and 2) reduced auditory thresholds. Both phenomena would lead to an increased auditory sensitivity. Moreover, the dynamic modulation of auditory sensitivity might be an important mechanism to provide protection from sensory bombardment inside the noisy swarm. It is even plausible that high RDL conductance at times of the day other than swarm time “silence” the auditory nerve, explaining why male attraction to female sounds are limited to the mating period. In any case, our data strongly suggest that altering RDL signalling in mosquitoes could radically alter their ability to detect mating partners. Given that RDL is an insecticide target, future research will explore the implications of our findings for mosquito control or insecticide management strategies.

**Figure 6:**
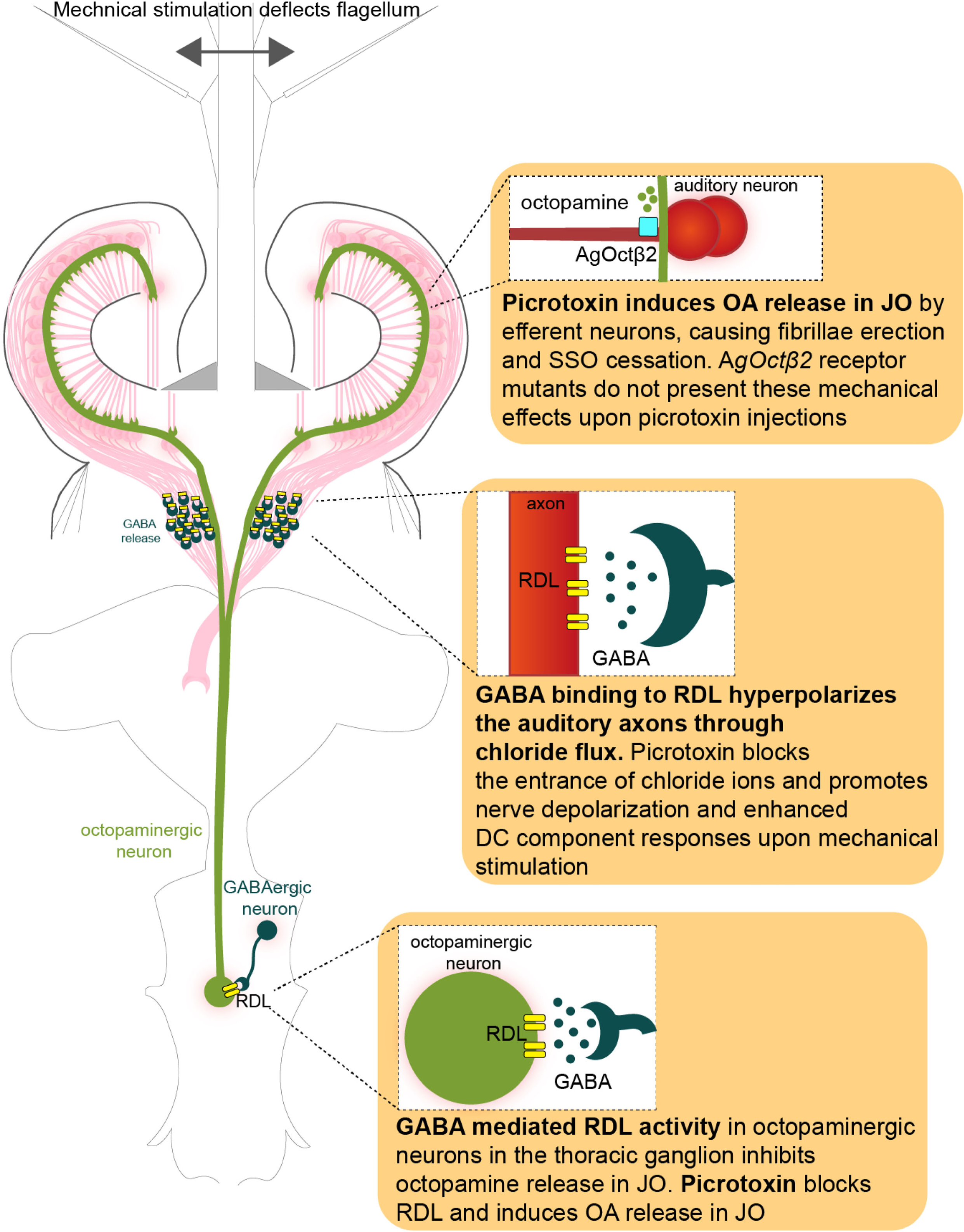
Proposed model for RDL auditory role in malaria mosquitoes. OA: octopamine, DC: direct current, JO: Johnston’s organ, SSO: self-sustained oscillations.

---

**Supplementary Table 1:** HCR probes were designed to target all splice isoforms of RDL (AGAP006028).

**Supplementary Figure 1: Picrotoxin effects on stimulated auditory responses after control injection.** Male mosquitoes were exposed to frequency-modulated sweep stimulation before and after control injection (Ringer 0.1 % DMSO). The data is displayed in the same way as in Fig. 4 and 5 (see legends for a detailed description of data shown). **A)** Envelopes of antennal responses were either fit with splines (top, white background), or a Damped Harmonic Oscillator (DHO) function (bottom, ochre background). See methods for details of fitting procedures. For splines (top, white background): left strip plots show the peak-to-peak amplitude of flagellar displacements between upper and lower splines. Data extracted from individuals before (black) and after injection (red). Lower right strip plots show the stimulus frequency at the peak response. For DHO fits (bottom, ochre background): left strip plots show the Q factor (i.e. tuning sharpness) extracted from the fits, and right strip plots show the oscillator (top) and peak frequency (bottom) – another set of parameters that can be extracted from DHO fits (see (6)). Control injection did not cause any changes in the auditory mechanics. **B)** AC component extracted from applying a highpass filter to compound action potential (CAP) responses. Left strip plots show spline-to-spline amplitude at the peak of the lower spline. Lower right strip plots show the stimulus frequency at the peak response. The frequency of the stimulus causing peak AC component responses did not change after control injection. **C)** DC component extracted from applying a lowpass filter to compound action potential (CAP) responses. All strip plots are the same as for high-pass filtered data. Control injection did not cause changes in DC component responses. DHO: damped harmonic oscillator. Sample size = 9, some samples had to be omitted, particularly after injection, as we were unable to perform robust spline-fitting on averaged data.

## Notes

https://github.com/mosquitome/sweep-n-sleep

### Competing Interest Statement

The authors have declared no competing interest.

