## Supplementary figures and images for "The GABA_A_ receptor RDL modulates the auditory sensitivity of malaria mosquitoes"

### Supplementary figure 1

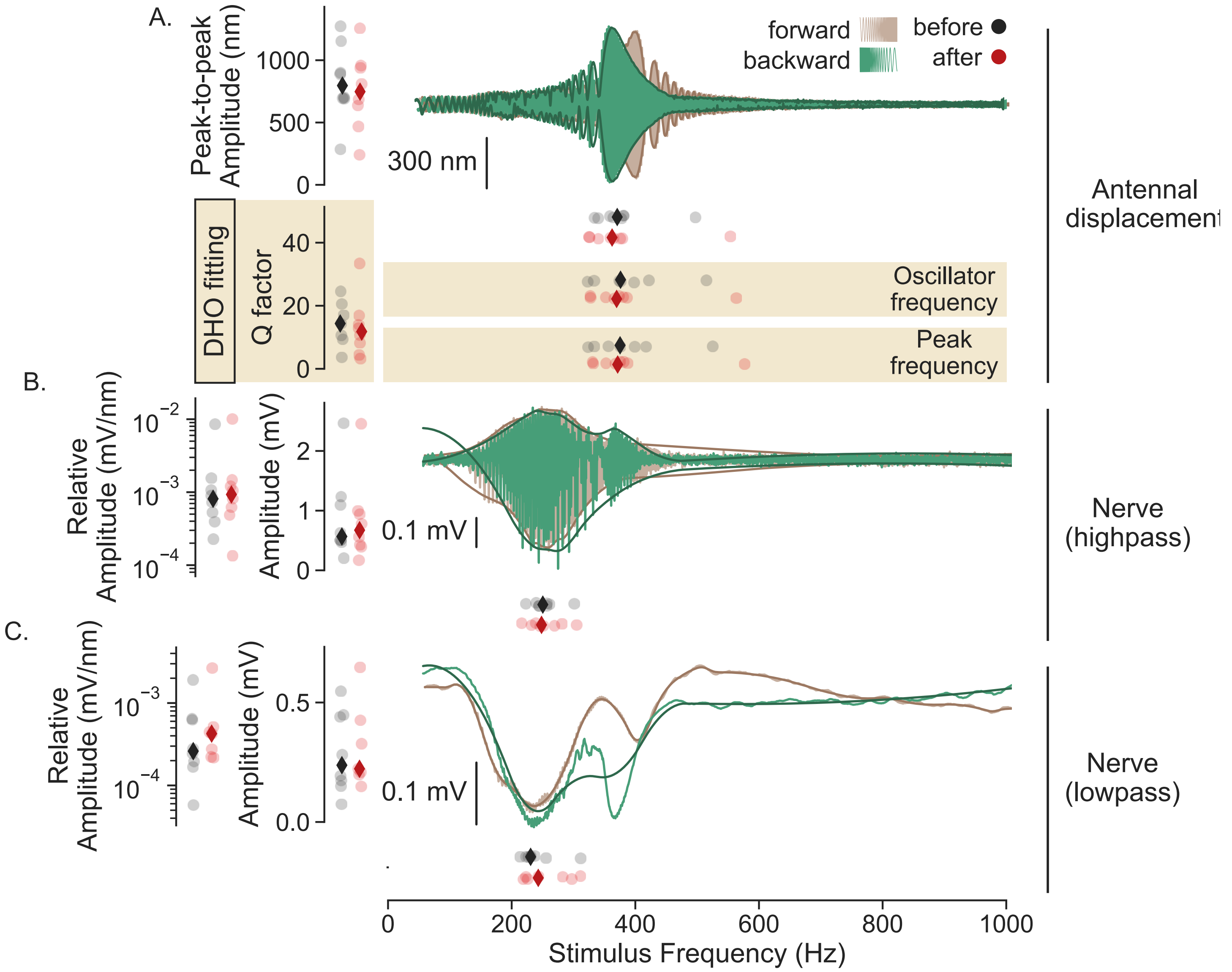
